# Encoding of hunger by the neuronal epigenome slows aging in *Drosophila*

**DOI:** 10.1101/2022.07.21.501022

**Authors:** KJ Weaver, RA Holt, E Henry, SD Pletcher

## Abstract

Hunger is, by necessity, an ancient motivational drive, yet the molecular nature of homeostatic pressures of this sort and how they modulate health and physiology are largely unknown. Here we show that the molecular encoding of hunger slows aging in *Drosophila*. We identify the branched-chain amino acids (BCAAs) as dietary hunger signals that extend lifespan despite increasing food intake when reduced, and in parallel show that optogenetic activation of a subset of hunger-promoting neurons is sufficient to recapitulate these effects. We find that remodeling of the neuronal histone acetylome is associated with dietary BCAA reduction, and that this requires BCAA metabolism in specific subsets of neurons. Preventing the histone acetylome from being molded by dietary BCAAs abrogates both increased feeding and extended lifespan. However, the mechanisms that promote feeding and modulate aging downstream of alterations in histone acetylation occur through spatially and temporally distinct responses; differential usage of the histone variant H3.3A in the brain is an acute response to hunger that promotes increased feeding without modulating lifespan, while a prolonged experience of hunger may slow aging by promoting a beneficial decrease of a set-point around which hunger levels are regulated. Identification of a molecular basis for the encoding of hunger and demonstration of its sufficiency in extending lifespan reveals that motivational states alone are deterministic drivers of aging and behavior.

## Introduction

The relationship between an animal and its diet influences behavior and aging in remarkable ways. The physiological need for nutrients motivates animals to forage and feed, and forced limitation in food availability slows aging across taxa(*1*). Both effects derive not only from the energetic content of the diet but also its composition(*2–4*). Many animals eat until they have consumed a specific amount of protein, for example, and the protein:carbohydrate ratio that results in part from seeking this target is a major factor in modulating lifespan(*5*). Remarkably, an animal need not consume its diet to be affected by it. In mice, appetite is promoted by environmental cues that predict future food consumption, presumably by influencing broader neural states that specify nutrient-specific drive, or hunger(*6, 7*). Similarly, the taste and smell of specific nutrients modulate lifespan in, among other species, the fruit fly, *Drosophila melanogaster*, and the nematode, *Caenorhabditis elegans*(*8–11*).

It seems likely that the effects of diet on aging and behavior share mechanistic foundations in the motivational states they promote, yet little is known about the molecular nature of these states. Many animals, including humans, develop a motivational drive for protein, which has also been described in *Drosophila* in response to starvation and mating(*12–14*). The existence of this drive and its ability to influence physiology is demonstrated by the observation that perception of protein-containing food without its consumption reverses the beneficial effects of protein-restricted diets in *Drosophila* and *C. elegans.* (*11, 15, 16*). Serotonin, together with neurons that express a specific serotonin receptor (5-HT2A), modulate this hunger state, and loss of 5-HT2A increases lifespan up to 50% in nutrient-rich conditions(*17*). Observations like these led us to consider the hypothesis that the neural states that encode the motivation to seek food and that define hunger *per se* may be sufficient to slow aging, independent of any changes in nutrient intake that may result.

Here we establish that hunger modulates aging in *Drosophila* and demonstrate that epigenomic encoding of this motivational state promotes feeding and modulates lifespan through partially distinct downstream mechanisms. We show that two models of hungry flies – by reducing dietary branched-chain amino acids or by activation of neurons that evoke hunger - are long-lived, despite consuming more calories and total protein. We establish that specific subsets of *Drosophila* neurons use BCAA metabolism to promote the decoration of histone tails with acetylation marks in the brain; a plasticity that is required to encode hunger in these models. Finally, we present evidence to suggest that prolonged hunger alters a set-point around which appetite is regulated and that this adaptation is an important component of a slowed aging process.

## Results

To investigate how neural states motivate feeding and modulate aging, we focused our investigations on the branched-chain amino acids (BCAAs) primarily because reducing them in the diets of mammals and flies increases protein appetite and extends lifespan(*18, 19*). We used a chemically defined diet to titrate and manipulate the BCAAs without altering other dietary components(*20*). We first designed a reference holidic diet (RD), around which we were able to manipulate BCAAs within a range of concentrations consistent with standard diets used for *Drosophila* aging studies that ensure against mal- and over-nutrition. We enforced equal concentrations of all other non-essential amino acids and non-BCAA essential amino acids across diets, which allowed us to study the effects of BCAAs without the confounding effects of general amino acid deficiency that are mediated by well-described mechanisms (Fig. 1A)(*13, 21–23*). This led to diets of modestly different caloricity. However, caloric content has been shown to be less impactful than dietary composition in modulating lifespan and subsequent experiments ruled out differences in calories as a cause of the dietary effects we observed(*2*).

**Figure 1.**
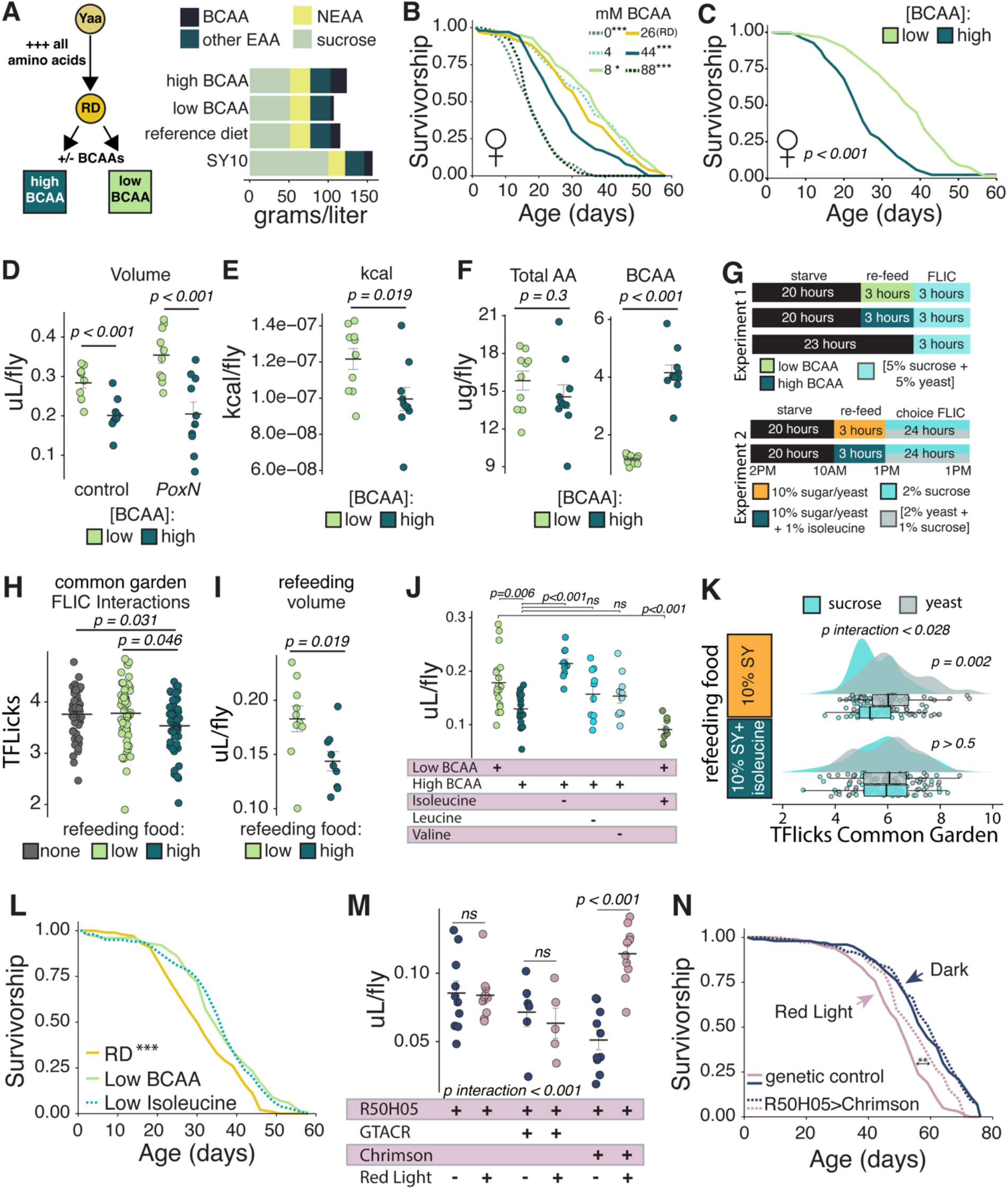
Hunger extends lifespan independent of appetite in dietary and genetic models of hungry flies. **(A)** Schematic (left) and relative composition (right) of experimental diets. Details in Supplementary Tables S1-2 and Methods. Yaa=baseline diet as in (*20*), RD=reference diet, containing 51.36g/L carbohydrate, 34.9g/L non-essential amino acids, 22.56g/L non-BCAA essential amino acids and, 12.39g/L BCAAs. **(B)** Lifespan of *Canton-S* flies on diets of indicated BCAA concentration (log-rank test, p-values derived by comparison to RD, N=159-172). **(C)** Lifespan of *Canton-S* flies on low- or high-BCAA diets (log-rank test, N=173-175). **(D-F)** 24-hr Con-Ex measurement of *w^1118^* or *PoxN* flies on indicated diets. Volume (D), kcal (E), and total or BCAA-only amino acid intake (F) (two-tailed Student’s t-test). **(G)** Schematic of FLIC experimental designs used in H-I (top panel) and K (bottom panel). **(H)** FLIC interactions in common garden (one-way ANOVA with Tukey’s post-hoc. Experimental replicates are pooled, N=52-55). **(I)** Con-Ex measurement of volumetric intake during re-feeding period (two-tailed Student’s t-test). **(J)** Con-Ex measurement on diets with individual BCAA reductions (one-way ANOVA with Tukey’s post-hoc). **(K)** FLIC interactions in food choice environment (2% sucrose or [2% yeast + 1% sucrose]) after re-feeding SY food +/- 1% isoleucine (two-way ANOVA with Tukey’s post-hoc. Experimental replicates are pooled, N=59-60). **(L)** Lifespan of *Canton-S* flies on RD, low-BCAA, or low-isoleucine diets (log-rank test, p-values derived by comparison to low-BCAA diet, N=171-177). **(M-N)** Con-Ex and lifespan measurements of flies carrying R50H05-GAL>UAS-CsChrimson or R50H05-GAL4/w-;CS controls exposed to red light for 12 hours per day or kept in constant darkness. (M) 24-hr Con-Ex measurement (two-way ANOVA with Tukey’s post-poc). (N) Lifespan measurement (log-rank test, N=101-123). All FLIC data are expressed as box-cox transformation to the 0.25 power (termed TFLicks) to achieve normality. *p<0.05, **p<0.01,***p<0.001.

Consistent with previous reports, dietary BCAA concentration modulated fly lifespan(*19, 24*) (Fig. 1B). Lifespan extension was larger in female flies and was maximized on a diet containing roughly 1/3^rd^ of the BCAA content of the reference diet, which is consistent with lifespan extension observed with dilution of the standard laboratory diet (Fig. 1B, Fig. 1C, Fig. S1A)(*2, 3*). We subsequently focused our investigation using female flies because of their known, heightened lifespan and neuronal responses to protein availability, which we also observed(*25, 26*). We chose two diets from the range of those initially tested, hereafter termed “low -BCAA” (8mM BCAAs/5.6% w/v total amino acid) and “high-BCAA” (44mM BCAAs/7.2% w/v total amino acid), to investigate in detail.

Dietary BCAAs also modulated food intake. Using a method that determines how much volume a group of flies consume in 24 hours by feeding them on blue-dyed food and then collecting their excretions (termed “Con-Ex” for Consumption-Excretion)(*27, 28*), we found that flies on low-BCAA diets consumed more food volume compared to those on high-BCAA diets (Fig. 1D, left panel). This was due to an increase in food intake by flies on low-BCAA diets rather than a decrease by flies on high-BCAA diets because food intake was also increased compared to flies on our RD (Fig. S1B). Differences were unlikely a result of food taste; *Pox-Neuro* flies who have extreme deficits in chemosensation also consumed more food on low-BCAA diets (Fig. 1D, right panel). Increased volume intake by flies on low-BCAA diets resulted in significantly higher caloric intake (Fig. 1E) but in similar amounts of total amino acids (ug/fly) eaten between flies fed low- or high-BCAA diets (Fig. 1F, left panel), while BCAA intake was significantly reduced (ug/fly) for flies fed the low-BCAA diet (Fig. 1F, right panel). Thus, flies fed our low-BCAA diet consumed more calories and similar amounts of total amino acids yet lived significantly longer.

### Dietary BCAAs influence hunger states

The relative increase in food intake we observed when flies live on low-BCAA diets led us to speculate that reducing dietary BCAAs created a food environment that promoted a heightened and persistent state of hunger. Measuring hunger can be challenging in a simple model system; animals eat for many reasons, and total food intake on homogenous mixtures of nutrients may not be as indicative of hunger levels as feeding paradigms that quantify precisely when and how often individual animals interact with specific food sources(*6, 13, 17, 29, 30*). We therefore devised a refeeding assay to determine how BCAAs influence hunger. In this paradigm, starved flies are refed a measured 3-hour bolus of test food, such as low- or high-BCAA, and are then placed into a common food environment (termed “common garden”) where we measure food interactions using the Fly Liquid Food Interaction Counter (FLIC)(*31*) (Fig. 1G, Experiment 1, top panel). We reasoned that if BCAAs did influence a state of hunger, then manipulating their concentration in the test food during refeeding would alter future food interactions during assessment in the common garden.

We observed that flies refed a low-BCAA diet subsequently interacted more often with the sucrose/yeast food in the common garden than did flies refed a high-BCAA diet and, in fact, interacted as frequently as did flies that were not refed at all (Fig. 1H). This was not due to a reduction in total calories or amino acids consumed during the refeeding period: flies ingested significantly more food volume and also more calories, as measured by Con-Ex, when refeeding on a diet of low-BCAAs than they did when refeeding on a high-BCAA diet (Fig. 1I), although total amino acid intake (ug/fly) was indistinguishable from flies refed high-BCAAs and BCAA consumption (ug/fly) was significantly less (Fig. S1C).

While the three BCAAs are commonly investigated together because they share similarities in their biochemical structures and functions, we next tested them individually for their role in promoting feeding. We measured food intake after reduction of each BCAA from the high-BCAA diet and found that decreasing isoleucine, but not valine or leucine or other pairwise combinations of amino acids, was required to increase feeding in the 24hr ConEx assay (Fig. 1J, Fig. S1D, S1E). Reducing dietary isoleucine increased the total calories consumed (Fig. S1E) and was the only dietary manipulation that we observed as sufficient to promote increased total amino acid intake (Fig. S1D). Furthermore, increasing isoleucine alone from our low-BCAA diet to match the concentration in our high-BCAA diet was sufficient to decrease food intake (Fig. 1J), demonstrating that isoleucine acts as a dietary signal to modulate feeding.

Having established that isoleucine was sufficient to modulate volumetric food intake, we next returned to our refeeding paradigm to determine how nutrient-specific drives may be affected by this amino acid using the FLIC to measure behavior in a food-choice environment. Choices between carbohydrate- or protein-rich food are thought to be tuned by increased protein demand in starved animals, who exhibit a robust increase in interactions with protein-rich food compared to carbohydrate-rich food, and to serve as a useful proxy for measuring protein-specific hunger(*31–33*). We reasoned that if isoleucine has a specific influence on protein hunger, then manipulating its concentration in the test food during refeeding would be expected to alter future choices between protein or carbohydrate food during the subsequent assessment period. For these experiments, we allowed flies a bolus of refeeding for three hours on either conventional 10% sugar-yeast food or the same food to which we added 1% isoleucine. We then measured over the next 24hrs the frequency of individual fly interactions with either carbohydrate- or protein-rich food in the choice environment (Fig. 1G, Experiment 2, bottom panel). Control flies that were refed conventional sugar-yeast food exhibited more interactions with protein-rich food compared to the carbohydrate-rich food, establishing that the three hours of refeeding before the test period were insufficient to fully satiate the animals and that they maintained a heightened level of protein drive (Fig. 1K, top). On the other hand, flies that were refed food with added isoleucine had statistically indistinguishable interactions with carbohydrate-compared to protein-rich food (Fig. 1K, bottom). Thus, levels of dietary isoleucine modulate protein-specific appetite, suggesting that our low-BCAA diet could promote a heightened state of hunger, perhaps by influencing protein drive.

### Hunger states influence lifespan

We next asked whether restriction of isoleucine alone was capable of increasing lifespan. We reduced all BCAAs or just isoleucine from our reference diet and found that isoleucine reduction (but not valine or threonine) was sufficient to extend lifespan to a similar degree as reducing all BCAAs, despite increased intake of calories, total amino acids, and carbohydrates (Fig. 1L, Fig. S1F). This observation is consistent with our conjecture that the motivational state of hunger itself, rather than the availability or energetic characteristics of the diet, might slow aging. Further support for this idea was provided by the observations that our diets did not have significant effects on egg laying, activity levels, triglyceride and protein levels, and activation of mTOR/AKT pathways (Figs. S2, A-E), which are commonly associated with manipulations that modulate lifespan through changes in nutrient availability or toxicity.

To determine whether hunger itself might slow aging, we sought a way to induce it independent of dietary manipulations and examine its effect on lifespan. Neurons whose activity evokes hunger in the fly have been described, but their role in modulating lifespan has not been examined(*34*). These neurons can be manipulated by using flies that express the GAL4 transgenic activator under the R50H05 driver (R50H05-GAL4). By targeting expression of the light-sensitive cation channel CsChrimson to these neurons, we created flies in which R50 hunger neurons were optogenetically activated at will to artificially generate hunger and to increase feeding when adult flies were exposed to red light. We observed that these flies consumed twice as much food as did flies of the same genotype that were kept in the dark or that expressed a green-light sensitive opsin channel (GTACR), consistent with prior reports (Fig. 1M). Remarkably, we also found that inducing a heightened state of hunger by activating these neurons for 12 hours each day during the light period significantly extended lifespan relative to a genetic control strain, which was not observed in the dark where flies are modestly long-lived in general – a known phenomenon that has been reported by our lab and others (Fig. 1N, Fig. S3). We interpreted this as a stark confirmation of our conjecture that hunger itself slows aging.

### Molecular encoding of hunger by the neuronal epigenome

We next turned to defining the molecular interactions among BCAAs, hunger neurons, and the brain to determine how hunger might be molecularly encoded. Our attention was drawn to the epigenome because alterations in histone proteins are involved in the generation of motivated behaviors, are a hallmark of aging, and are regulated by nutrients(*35–38*). Furthermore, histone acetylation is especially sensitive to the availability of dietary nutrients that generate acetyl-CoA, and BCAAs and isoleucine specifically are metabolically fated to produce this substrate(*39, 40*).

We first examined whether dietary BCAAs influenced relevant histone post-translational modifications. We found that histone H3K9 acetylation was significantly decreased in fly heads within one week of feeding a low-BCAA diet (Fig. 2A). Subsequent experiments revealed that H3K27 acetylation, but not H3K9 methylation, was also reduced when flies were fed low-BCAA diets. All significant effects of dietary BCAAs on histone PTMs were abrogated by feeding flies the histone deacetylase inhibitor, sodium butyrate (Fig. 2C). We also observed an unexpected decrease in total histone H3 abundance in heads and brains, although H3 mRNA was increased (Fig. 2A-C, Fig. S4). The significant decrease in total histone H3 was independent of changes in histone H4, which were statistically non-significant between diets (Fig. 2D). Inducing hunger independent of diet by activation of R50 hunger neurons also reduced histone H3 abundance in fly heads (Fig. 2E), and reducing isoleucine alone, which was important for modulating feeding and lifespan (e.g., Fig. 1), partially recapitulated the effects of BCAAs on histone H3 abundance, suggesting that epigenetic changes may be causally linked to one or both hunger phenotypes (Fig. 2D).

**Figure 2.**
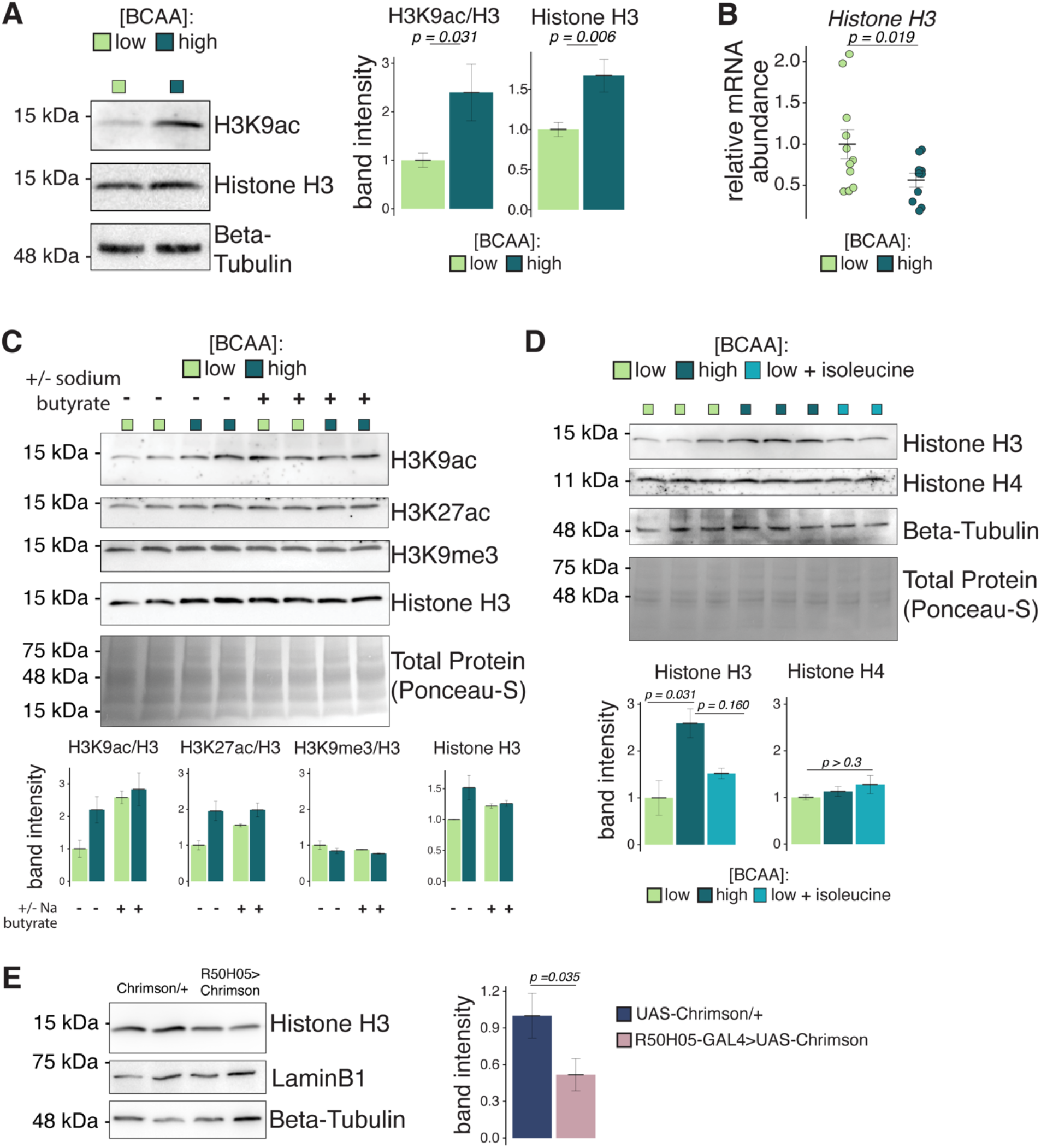
Dietary BCAAs reshape the neuronal epigenome by altering histone acetylation and histone H3 abundance. **(A)** Representative western blot for H3K9ac and total histone H3 in *Canton-S* fly heads after exposure to indicated diets for 5-7 days, quantified in right panel (one-way ANOVA. 10 heads per biological replicate, experimental replicates are pooled, N=11). **(B)** RT-qPCR measurement of relative mRNA abundance in *Canton-S* heads after 5-7 days on low- or high-BCAA, values are normalized to low-BCAA treatment (one-way ANOVA. 10 heads per biological replicate, experimental replicates are pooled, N=10-11). **(C)** Western blot for histone PTMs in *Canton-S* fly heads after exposure to indicated diets +/- 100 mM sodium butyrate for 5-7 days, bands are quantified in bottom panel and normalized to low-BCAA treatment (10 heads per biological replicate, N=2). **(D)** Western blot for histone H3 and H4 in *Canton-S* heads on low-BCAA, high-BCAA, or [low-BCAA + high isoleucine] diets, quantified in bottom panel and normalized to low-BCAA treatment (one-way ANOVA with Tukey’s post-hoc. 10 heads per biological replicate, N=2-3). **(E)** Western blot for histone H3 in heads of flies carrying R50H05-GAL>UAS-CsChrimson or UAS-CsChrimson/w-;CS controls exposed to red light for 12 hours, quantified in right panel and normalized to UAS-CsChrimson/w-;CS control (one-way ANOVA. 10 heads per biological replicate, N=4-5).

We next focused on the unexpected finding that histone H3 abundance was reduced by low-BCAA diets. Increased H3 mRNA levels coupled with decreased protein abundance led us to hypothesize that BCAAs may influence turnover of histone H3, such that it may happen more quickly on low-BCAA diets. In the brain, histone H3 is marked for eviction from chromatin by acetylation marks and can be replaced by the histone variant H3.3(*41, 42*). This variant accumulates with age and is thought to decorate regions of actively transcribed chromatin(*43*). We found that it is also transcriptionally increased in the heads of our flies fed low-BCAA diets (Fig. 3A). To visualize the incorporation and removal of histone H3.3 into neuronal chromatin, we used an inducible pan-neuronal driver (Nsyb-GeneSwitch-GAL4) to pulse and then chase fluorescently tagged H3.3 into the brain(*41*). We observed higher H3.3-GFP signal in fly brains after one week of feeding on low- vs. high-BCAA diets, indicating an increase in its stability or persistence in chromatin (Fig. 3B, lower panel; quantified in Fig. 3D), which was not due to changes in initial H3.3-GFP incorporation (Fig. 3B, top panel; quantified in Fig. 3C) or to differences in protein degradation more generally (Fig. 3D). These data indicate differential usage of histone H3 and the variant H3.3A in the fly brain following feeding on a low-BCAA diet.

**Figure 3.**
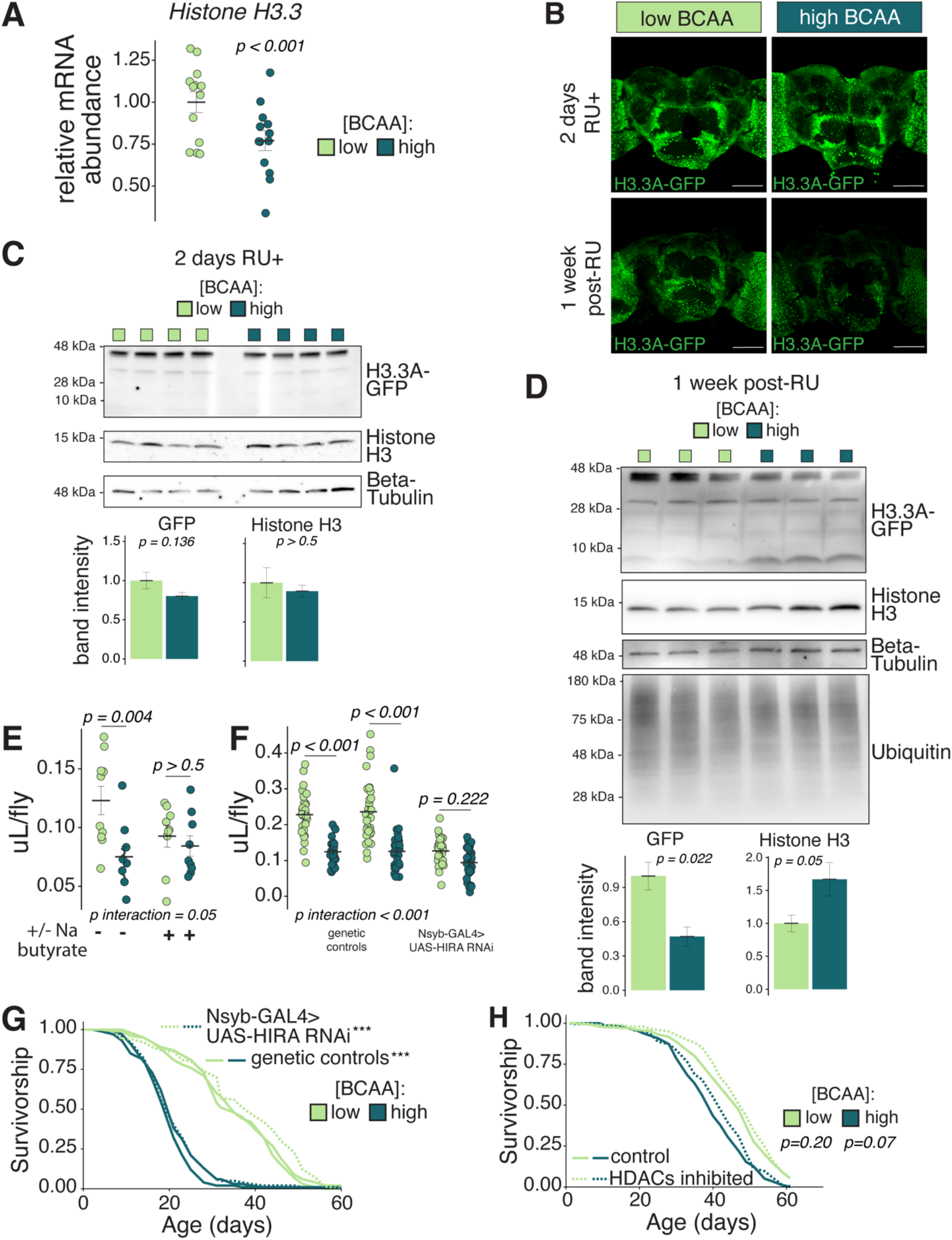
Utilization of the histone variant H3.3A in the brain is modulated by BCAAs to stimulate feeding but is not required to extend lifespan. **(A)** RT-qPCR quantification of relative mRNA abundance of *H3.3A* in *Canton-S* heads after 5-7 days on low- vs. high-BCAA diet, values are normalized to low-BCAA treatment (one-way ANOVA. 10 heads per biological replicate, experimental replicates are pooled). **(B-D)** Pulse-chase of fluorescently labelled H3.3A in brains (B) or heads (C-D) of flies carrying *Nsyb-GeneSwitch-GAL4>H3.3A-GFP* on low- or high-BCAA diets. (B) Representative confocal images of brains immediately following the 2-day pulse of H3.3A-GFP induced by BCAA food + RU486 (top panel) and 1 week after the pulse (bottom panel). Scale bar 100μm. (C) Western blot of H3.3A-GFP (predicted molecular weight = 42 kDa) and histone H3 immediately following the 2-day pulse of H3.3A-GFP induced by BCAA food + RU486, quantified in bottom panel (one-way ANOVA, 10 heads per biological replicate). (D) Western blot of H3.3A-GFP (one-way ANOVA), histone H3 (one-tailed Student’s t-test), and ubiquitin (one-way ANOVA, p=0.571) 1 week after the H3.3A-GFP pulse, quantified in bottom panel (10 heads per biological replicate). **(E)** Con-Ex measurement of 24-hr food intake after 5 days on BCAA diets +/- 100 mM sodium butyrate (two-way ANOVA with Tukey’s post-hoc). **(F)** Con-Ex measurement of 24-hr food intake after 5-7 days on BCAA diets from flies carrying *Nsyb-GAL4>UAS-HIRA RNAi* or controls (*Nsyb-GAL4/w-;CS* and *UAS-HIRA RNAi/w-;*CS) (two-way ANOVA with Tukey’s post-hoc, experiment replicates are pooled). **(G)** Lifespans of flies carrying *Nsyb-GAL4>UAS-HIRA RNAi* or controls (*Nsyb-GAL4/w-;CS* and *UAS-HIRA RNAi/w-;CS*) (log-rank test, N=184-202). **(H)** Lifespan of flies on BCAA diets +/- 100 mm sodium butyrate (log-rank test, N=186-196).

To determine whether the observed differences in histone acetylation or variant usage were required for feeding or lifespan differences between the diets, we began with a pharmacological approach to broadly inhibit histone deacetylases (HDACs), which prevented diet-dependent differences in histone acetylation (Fig. 2C). Feeding flies the HDAC inhibitor sodium butyrate for one week abrogated increased feeding on low-BCAA diets, indicating that the histone acetylome could either encode hunger itself or be a permissive response to hunger that modulates feeding (Fig. 3E). To distinguish between these possibilities, we fed flies HDAC inhibitors for life and found that this extended the lifespan of flies on high-BCAA food. Dietary modulation of histone acetylation therefore regulates both feeding and lifespan, supporting the idea that the histone acetylome encodes hunger rather than feeding *per se* (Fig. 3H). We also tested whether incorporation of histone variant H3.3 was required for the effects of hunger on feeding and lifespan by using pan-neuronally expressed RNAi to knock-down the protein chaperone HIRA, which is required for the exchange of H3 for H3.3(*44*). We observed that this also abolished increased feeding on a low-BCAA diet (Fig. 3F), but surprisingly, it had no effect on lifespan (Fig. 3G). Histone H3/H3.3 swapping is therefore required for modulating feeding in response to hunger but is dispensable for hunger’s effects on lifespan.

### Metabolic intermediates link BCAAs to feeding and lifespan

How might BCAAs from the diet promote modifications to the neuronal epigenome? The BCAAs are unusual amino acids because, in mammalian systems, they bypass metabolism in the liver and are instead metabolized in target tissues by the enzymes branched-chain aminotransferase (BCAT) and branched-chain alpha-ketoacid dehydrogenase (BCKDH)(*40, 45, 46*). We found that *BCAT* mRNA abundance was reduced in the heads of flies fed low-BCAA diets (Fig. 4A). Knocking-down *BCAT*, a homolog of mammalian *BCAT2*, with the pan-neuronal Nsyb-GAL4 driver reduced histone H3 abundance to statistically indistinguishable levels on low-compared to high-BCAA diets (Fig. 4B). We consulted the Fly Cell Atlas and found that *BCAT* is expressed in approximately 80 neurons that co-express the promoter used to label R50 hunger neurons, *SerT* (*47*) (Fig. S5). Strikingly, BCAT knock-down only in the R50 hunger neurons also resulted in statistically indistinguishable levels of histone H3 abundance and reduced H3K9ac on low-compared to high-BCAA diets (Fig. 4C). It also strongly reduced diet-dependent differences in lifespan by significantly extending the lifespan of flies on high-BCAA diet (Fig. 4D). This is unlikely to be a result of deficiencies in neuronal activity upon knockdown of *BCAT* because optogenetic inhibition of the R50 hunger neurons had statistically non-significant effects on the lifespan of flies fed high-BCAA diets (Fig. S6). These findings indicate that R50 hunger neurons use BCAA metabolites to modulate aging via a mechanism that does not require neuronal depolarization. We were surprised, however, to observe that *BCAT* knockdown in hunger neurons did not abrogate feeding differences and may have in fact exacerbated them (Fig. 4E). These results suggest complexity in the effector pathways that increase feeding and modulate aging in response to hunger and reinforce the notion that they are, at least in part, anatomically distinct.

**Figure 4.**
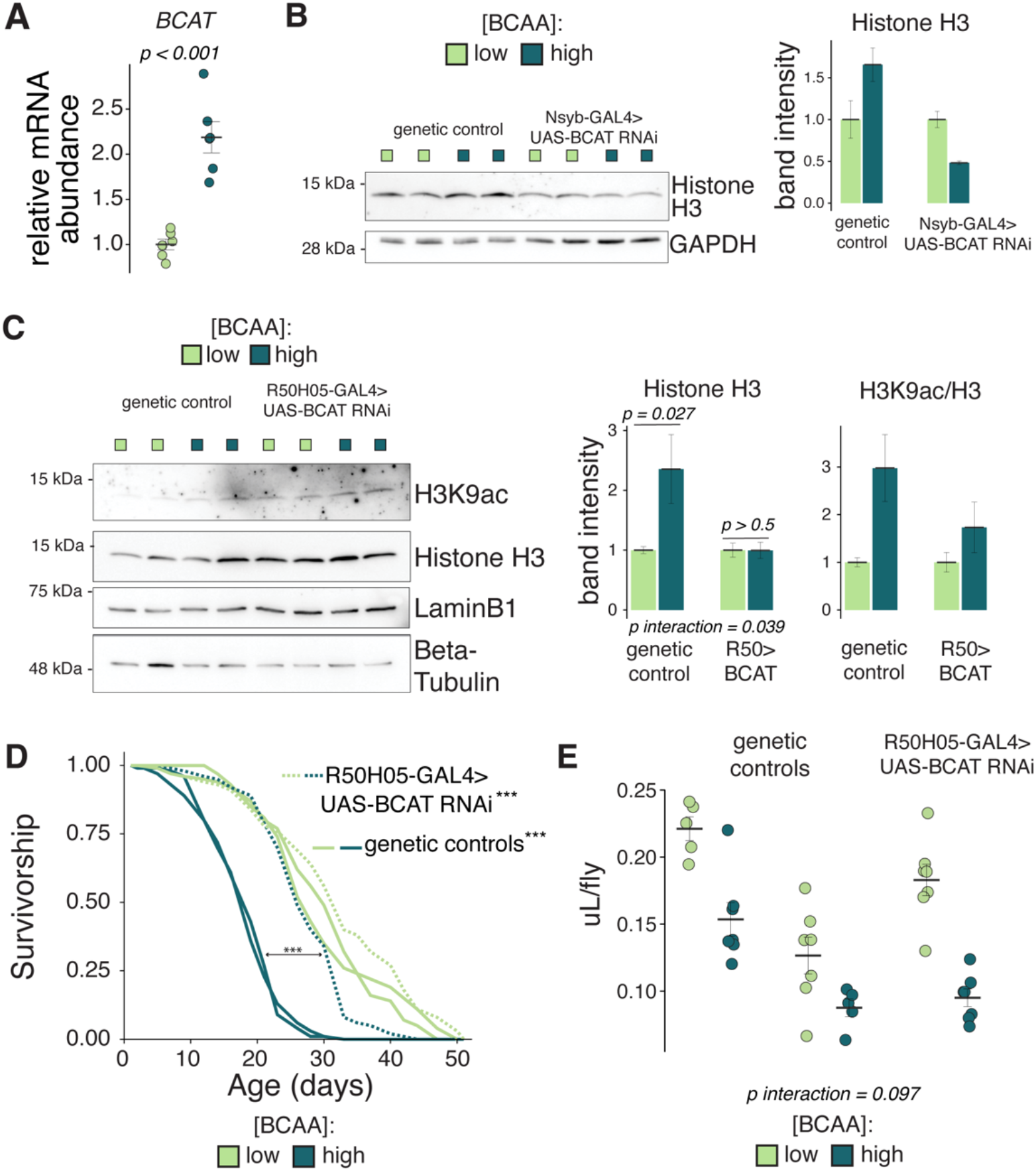
Hunger neurons use BCAA metabolism to regulate histone abundance and modulate lifespan independent of feeding. **(A)** RT-qPCR quantification of relative mRNA abundance of *BCAT* (*CG1673)* in *Canton-S* heads after 5-7 days on BCAA diets, values are normalized to low-BCAA treatment (one-way ANOVA, 10 heads per biological replicate). **(B)** Western blot of histone H3 in heads of flies carrying *Nsyb-GAL>UAS-BCAT RNAi* or control (*Nsyb-GAL4/w-;CS)*, quantified in right panel and normalized to low-BCAA treatment (two-way ANOVA with Tukey’s post-hoc, N=2). **(C)** Western blot of histone H3 and H3K9ac in heads of flies carrying *R50H05-GAL4>UAS-BCAT RNAi* or control (*R50H05-GAL4/w-;CS)*, quantified in right panel (two-way ANOVA with Tukey’s post-hoc, experimental replicates pooled, N=5). **(D)** Lifespans of flies carrying *R50H05-GAL4>UAS-BCAT RNAi* or controls (*R50H05-GAL4/w-;CS* or *UAS-BCAT RNAi/w-;CS)* on low- or high-BCAA diets (log-rank test, N=91-100, ***p<0.001,). **(E)** Con-Ex measurement of 24-hr food intake after 5 days on BCAA diets in flies carrying *R50H05-GAL4>UAS-BCAT RNAi* or controls (*R50H05-GAL4/w-;CS* or *UAS-BCAT RNAi/w-;CS)* (two-way ANOVA).

### Considerations on the divergence of mechanisms linking hunger with feeding and aging

A subset of the R50 hunger neurons produce the neuromodulator serotonin, and we observed that expression of the genes *Trh*, which encodes tryptophan hydroxylase that functions as the first and rate-limiting step in serotonin synthesis, and *Tph/Henna*, which encodes tryptophan phenylalanine hydroxylase, were significantly increased in the heads of flies that were BCAA-restricted (Fig. 5A). We also observed increased antibody staining of serotonin itself in the cell bodies of the serotonergic PLP cluster, a subset of the R50 hunger neurons chosen for quantification due to their easily accessible and recognizable anatomical location in the fly brain (Fig. 5B-C, quantified in 5D). Given that inhibition of BCAT in the R50 neurons seemed to exacerbate the feeding differences on our BCAA diets and that serotonin is known to both promote and suppress feeding depending on which serotonergic neurons are manipulated(*34*), we asked whether *BCAT* knockdown in the serotonergic network more broadly may produce opposite effects on feeding. We used the Trh-GAL4 driver, which is putatively expressed in all serotonergic neurons of the CNS, to target knock-down of *BCAT* and therefore BCAA metabolism in these cells. This manipulation prevented increased feeding on low-BCAA diets, suggesting that BCAA metabolites are used within distinct populations of serotonergic neurons to regulate feeding and aging (Fig. 5E).

**Figure 5.**
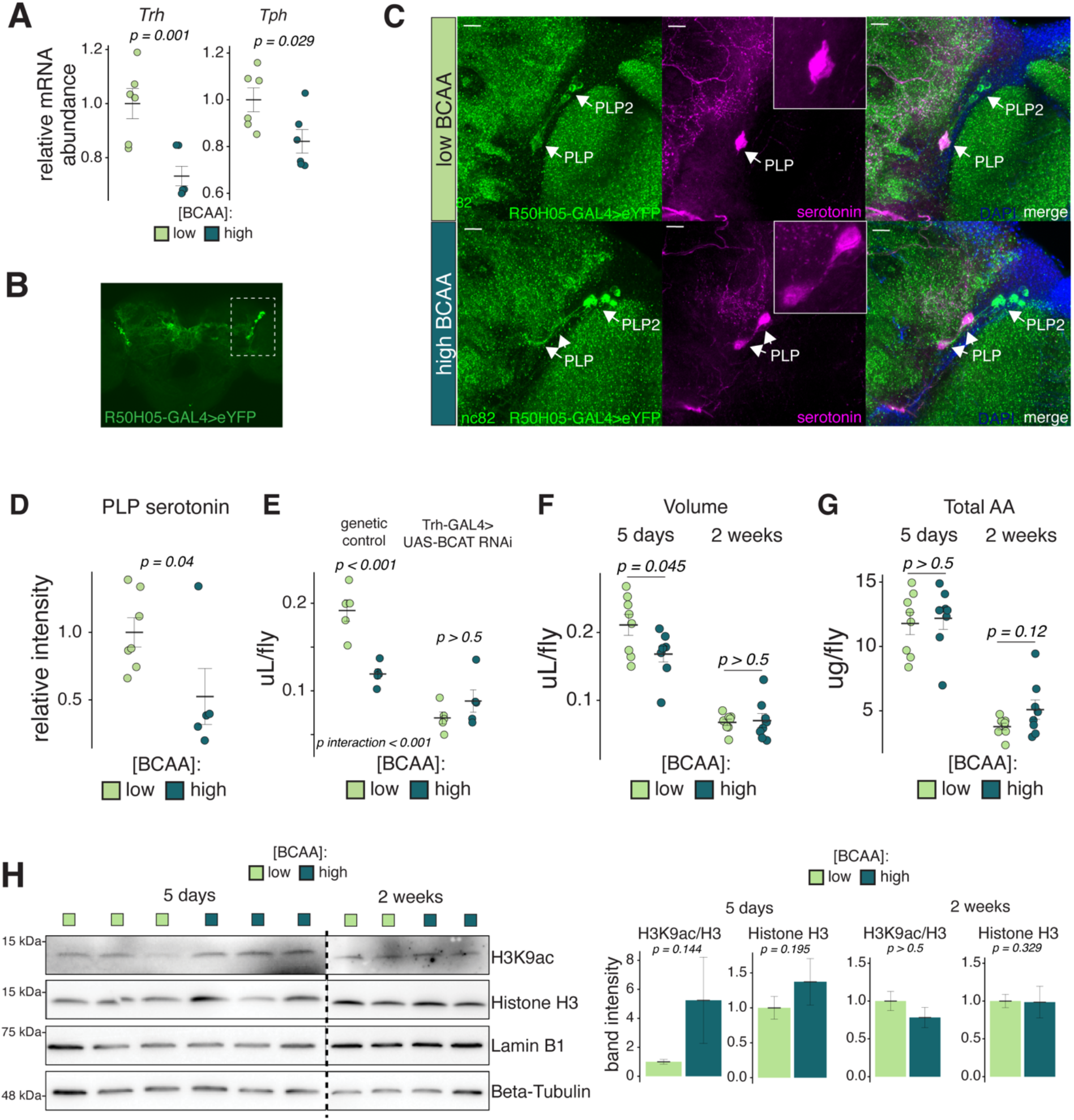
Spatially and temporally distinct processes link hunger to feeding and aging. **(A)** RT-qPCR quantification of relative mRNA abundance of *Trh* and *Tph* in *Canton-S* heads after 5-7 days on BCAA diets, values are normalized to low-BCAA treatment (one-way ANOVA. 10 heads per biological replicate). **(B-D)** Immunostaining for serotonin in PLP neuron cell-bodies in flies carrying *R50H05-GAL4>UAS-GTACR.eYFP* after 5-7 days on BCAA diets. (B) Maximum intensity projection of ten 1.5 μm stacks highlighting the PLP neurons (white box) in the posterior fly brain. Scale bar 100μm. (C) Representative confocal images of immunostaining for serotonin in PLP neurons of flies carrying *R50H05-GAL4>UAS-GTACR.eYFP (*green=nc82, magenta=serotonin, blue=DAPI, scale bar=10μm). Images are maximum intensity projections of ten 1.5 μm stacks through the PLP neurons (D) Quantification of (C) as described in Methods (one-way ANOVA, experimental replicates are pooled). **(E)** Con-Ex measurement of 24-hr food intake after 5 days on BCAA diets in flies carrying *Trh-GAL4>UAS-BCAT RNAi* or control (*Trh-GAL4/w-;CS)* (two-way ANOVA). **(F-G)** Con-Ex measurement of 24-hr food intake after 5 days or 2 weeks on BCAA diets in *Canton-S* flies; volume (F) and total amino acid intake (G) (two-tailed Student’s t-test). (H) Western blot of histone H3 and H3K9ac in heads of *Canton-S* flies after 5 days vs. 2 weeks on BCAA diets, quantified in right panel (one-tailed Student’s t-tests, 10 heads per biological replicate, experimental replicates are pooled, N=3-5).

Intrigued by the possibility that hunger effects on feeding and aging may be regulated by partially distinct pathways, we questioned if some of this complexity might also result from the time scale on which they occur. While we consider lifespan to be an outcome of chronic dietary effects, measurements of feeding quantify more acute responses that may or may not be stable over longer time periods. We observed that the increased feeding on low-BCAA food that was present after five days on the diet had receded after two weeks (Fig. 5F), resulting in reduced amino acid intake overall (Fig. 5G). This result was mirrored by temporal changes in the epigenome: patterns of changes in brain histone acetylation between diets after two weeks were reversed or eliminated when compared with similar measures after only five days (Fig. 5H). Thus, flies on low-BCAA diets presumably adapt to their food environment in a way that decreases the amount of protein they require, which may have beneficial consequences for aging (see model, Fig. S7).

## Discussion

Here we present behavioral and molecular evidence in favor of the idea that the neural state of hunger modulates aging. We show that this motivational state is molecularly encoded by the neuronal epigenome, and that increased feeding and extended lifespan are consequences that seem to be orchestrated by unique downstream responses to hunger. We found that the mechanisms that increase feeding and extend lifespan in response to low-BCAAs share the use of a BCAA-metabolism to histone acetylation pathway but diverge downstream of this programming. Our findings that dietary BCAAs modulated use of the histone variant H3.3; that this was required for increasing feeding, but dispensable for increasing lifespan; and that unique populations of serotonergic neurons require BCAA metabolism to modulate either feeding or aging, suggest that different neurons use different strategies to generate responses to hunger. Such complexity is consistent with emerging reports of interactions between metabolism and the epigenome that are driven in part by specialized enzymes that reside in distinct cellular compartments and in some cases distinct nuclear sub-compartments(*48–50*). In *C. elegans,* H3.3 is important for engaging transcriptional networks that underlie neuronal plasticity in response to environmental manipulations(*51, 52*). Perhaps neuronal plasticity is one strategy used by circuits that promote feeding to generate behavioral responses to hunger. Notably, protein appetite was previously shown to require plastic changes in a dopaminergic circuit, although a role for histone variant swapping in this process has not been determined(*33*). Thus, the cell-type specificity of the usage of histone variant H3.3 and the genomic loci to which it is targeted in response to dietary BCAAs remain unknown.

The observation that BCAA reduction acutely increased appetite but that this eventually subsided indicates that hunger may act as an allostatic stressor that, like other model homeostatic systems, acutely increases feeding while chronic hunger may promote physiological changes that lower a set-point around which appetite is regulated(*53, 54*). Perhaps it is this adaptative response, mediated by modifications to the epigenome in discrete neural circuits, that slows aging? This idea is consistent with several dietary manipulations that are known to extend lifespan: low protein diets (and methionine restriction in particular) generate protein-specific appetites and intermittent fasting paradigms putatively increase the frequency with which an animal experiences hunger without affecting total caloric intake(*5, 55–59*).

Others have reported that the effects of BCAAs on feeding and lifespan are non-specific in *Drosophila* and that such effects can be observed by restriction of other amino acids, which we did not observe(*19*). These differences are likely due to experimental design; Juricic et al. used holidic diets that were comparatively low in their overall amino acid content, which resulted in reduced activation of TOR signaling. We chose diets that were amino acid replete and had no effect on TOR activity, which revealed BCAA-specific effects, suggesting that interactions between total amino acid abundance and reductions of individual amino acids warrant further investigation.

Finally, our data support a role for chromatin reorganization as a link among hunger, feeding, and aging, but its influence on transcriptional states in individual neurons remains an exciting area to be explored. In the brain, transcriptional changes occur in response to dietary manipulations, and yet, analysis of single “longevity genes”, of which the list is continually growing, is typically insufficient to generalize about how diet modulates aging. This prompts us to speculate that the effects could require an overhaul of transcriptional programming more broadly(*60–62*). Chromatin accessibility patterns, gene expression profiles, and intracellular signaling pathways converge to determine transcriptional states which, although they are most often studied in the context of neuronal-*fate* determination, may also tune neuronal *states* in response to dietary or other environmental manipulations(*63, 64*). Future work that explores how specific nutrients, and their interactions, interact with each of these processes to shape the transcriptional environment of distinct neurons presents an important next step towards uniting seemingly disparate effects of diet on physiology and will likely provide new insight into how motivational states influence aging.

## Materials and Methods

### Fly stocks and husbandry

Fly stocks were maintained on a standard cornmeal-based larval growth medium and in a controlled environment (21C, 60% humidity) with a 12:12hr light:dark cycle. We controlled the developmental larval density by manually aliquoting 32 ul of collected eggs into individual bottles containing 25-50mL of food at 25C. Following eclosion, mixed sex flies were kept on SY10% medium (10% w/v sucrose and 10% w/v yeast) for 2-4 days until they were sorted by sex and transferred onto holidic food or kept on SY10% food, as needed, for experiments. Experimental flies were flipped to fresh food e/o day until completion of the experiment. Unless otherwise noted, we used mated female flies that were between 7-12 days old for all experiments. The following stocks were used for experiments: *Canton-S and w^1118^* were obtained from the Bloomington *Drosophila* Stock Center. *UAS-CsChrimson* (BDSC #55135 and #55136), *UAS-H3.3A-GFP* (BDSC #68241), *UAS-HIRA RNAi* (BDSC #35346), *GMR50H05-GAL4* (BDSC #38764), *Trh-GAL4* (BDSC #38388), *UAS-BCAT RNAi* (VDRC KK110229) were purchased from BDSC or the Vienna *Drosophila* Resource Center, as indicated. *PoxN* mutants were provided by J. Alcedo (Wayne State University). *Nsyb-GAL4* was provided by L. Buttitta (University of Michigan, MI) and Nsyb-GeneSwitch-GAL4 was obtained from A. Sehgal\ (Perelman School of Medicine, PA). *UAS-GTACR* was provided by M. Dus (University of Michigan, MI). All transgenic lines used in this study, with the exception of *UAS-H3.3A-GFP, UAS-HIRA RNAi, Nsyb-GAL4, and Nsyb-GeneSwitch-GAL4* were back-crossed 10 generations to *w-;Canton-S* prior to experiments.

### Holidic food

Holidic media were prepared according to previous protocols with some modifications^20^. For all experiments, we used the Yaa ratio of amino acids, but increased each amino acid, sucrose, and agar (see Extended Data Table 1). Briefly, agar, sucrose, branched-chain amino acids and amino acids with low solubility (L-Leucine, L-Isolecuine, L-Valine, L-Tyrosine) were added to solutions containing metal ions and cholesterol. The mixtures were autoclaved for 20 minutes. Filter-sterilized acetate buffer and solutions of the remaining amino acids, vitamins, nucelotides, inositol, choline, and preservatives were added while mixtures were stirred on a hot plate not to exceed 65C. After mixing, 2mL of food was dispensed into vials and stored at 4C until use, but no longer than 3 weeks.

### Drug administration

Sodium Butyrate was purchased from abcam (ab120948) and Trichostatin A was purchased from Cayman (Item No. 89730). Prior to the lifespan experiment, TSA was dissolved in 100% ethanol to a concentration of 1 mM, aliquoted, and frozen at −20°C. Food was prepared fresh e/o day as follows: a TSA aliquot was thawed and sodium butyrate was added as powder to the TSA stock. Appropriate holidic diet food was melted and allowed to cool slightly, then the inhibitor cocktail was added (final concentration of TSA was 10 μm and sodium butyrate was 10 mm), gently mixed, and dispensed to new vials. For Con-Ex or western blot experiments using only sodium butyrate, holidic food was melted and sodium butyrate was added as powder to a final concentration of 100 mm. Roughly 250 ul of sodium butyrate-containing food was layered on top of appropriate holidic food vials and cooled before use.

For the pulse-chase GeneSwitch experiment, RU486 (mifepristone) was dissolved in 80% (v/v) ethanol at 10 mM concentration and marked by blue dye (5% [w/v] FD&C Blue No. 1; Spectrum Chemical) and stored at −20°C. Holidic food + RU486 was prepared by melting appropriate holidic foods and adding RU486 to a final concentration of 200 μM.

### Lifespan measurements

Lifespans were measured using established protocols^65^. We established 6-10 replicate vials for each treatment, with 20 flies per vial. Flies were transferred to fresh media every 2-3 days, at which time dead flies were removed and recorded using the DLife system ^65^. Flies were kept in constant temperature (25C) and humidity (60%) conditions with a 12:12hr light:dark cycle. Normally, we conducted at least two experimental replicates of each lifespan experiment.

### Optogenetic Assays

Flies expressing UAS-CsChrimson in desired neuronal populations were reared as described above. Upon eclosion, females were sorted to SY10% medium containing 800 μm all-*trans*-retinal (from a stock solution of 100mM ATR in 100% ethanol) and kept in the dark for 2 days. Flies were then flipped to appropriate holidic diet food containing 400 μm ATR (and flipped to fresh ATR-containing food e/o day for the duration of the experiment) and moved to a custom rig containing 627nm (red) LEDs or 530nm (green) LEDs (Luxeon) or kept in the dark in the same incubator (Fig. S3). The custom rig is fully enclosed to prevent leakage of light and houses 48 individual vials surrounded by mirrors. Custom hardware and firmware were designed to allow the experimenter to control the LED intensity and a range of other light stimulus parameters. Lifespan and Con-Ex experiments used a stimulus frequency of 40 Hz and a pulse width of 800 ms. Flies were exposed to this protocol every day for 12 hours per day during the light period, followed by 12 hours of darkness.

### Consumption-excretion feeding assays

Con-Ex experiments were carried out as previously described^27,28^. Experimental female flies were sorted to appropriate BCAA diets (10 flies per vial, 8-10 replicates for each treatment) when they were >4 days old. Food was changed every 2-3 days until the appropriate experimental time-point was reached (5 days for standard experiments, unless otherwise indicated). After the dietary pre-treatment period, blue test food was prepared in removable caps by adding 1% (w/v) FD&C Blue No. 1 to the appropriate diets. Flies were moved to fresh vials with the removable blue-food caps on the top of the vials and were allowed to feed and excrete for 24 hours. Caps and flies were removed after the 24-hour test period and flies were counted. Excreted dye was collected by vortexing each vial with 3mL of water. Concentration of the dye was determined by absorbance at 630nm and compared to a standard curve of known concentrations. For short-term Con-Ex experiments lasting <3 hours, both excretions and flies were collected and measured. To measure internal blue concentration, groups of flies were homogenized in 1mL of phosphate-buffered saline containing 0.1% Trixon X-100 (IBI Scientific) for 30s at 30Hz using a QIAGEN TissueLyser. Concentration of internal extracts were determined by absorbance at 630nm and both internal and excreted concentrations were summed to determine total consumption.

### Fly Liquid-Food Interaction Counter (FLIC) assays

Flies were tested on the Fly Liquid-Food Interaction Counter (FLIC) system as previously described^31^(Sable Systems International). Female flies were starved for 20-24 hours in vials containing a kimwipe with 2mL of Milli-Q water. For single-choice FLIC experiments, flies were flipped to low-BCAA, high-BCAA, or fresh starvation vials at 10AM the morning of testing and allowed to feed for 3 hours. Re-feeding food was spiked with blue dye to ensure feeding, and flies with visually blue bellies were selected for the experiment. FLIC *Drosophila* Feeding Monitors (DFMs, Sable Systems International, model DFMV3) were loaded with a food solution containing 5% sucrose and 5% yeast extract (w/v) in 4 mg/l MgCl^2^. After re-feeding, flies were briefly anesthetized on ice and aspirated into the DFM chambers. We began recording immediately after loading flies (generally, loading all DFMs requires <10 minutes) and measured FLIC interactions for 3 hours. For FLIC experiments that contained a choice environment in the common garden, re-feeding food was prepared fresh for each experiment by melting SY10% medium with or without 1% L-Isoleucine (w/v). FLIC DFMs were loaded with food solutions containing either 2% sucrose OR [2% yeast extract + 1% sucrose] in 4 mg/l MgCl^2^ and FLIC interactions were recorded for 24 hours. Each DFM was loaded with flies from at least two treatment groups to reduce technical bias from individual DFM signals. FLIC data were analyzed using custom R code, which is available at https://github.com/PletcherLab/FLIC_R_Code. Default thresholds were used for analysis except for the following: minimum feeding threshold = 10, tasting threshold = (0, 10). Animals that did not participate (i.e. returned zero values) were excluded from analysis. FLIC data are generally non-normal, and thus are expressed as a box-cox transformation to the 0.25 power of the total interactions, which yielded normal distributions.

### Western Blots

Experimental flies were flash frozen in liquid nitrogen after appropriate diet or light exposures. Heads and bodies were separated using a metal sieve on dry ice, and 10 heads were pooled for each biological replicate. Heads were first pulverized to a fine powder using a plastic pestle on dry ice. Protein extraction was carried out on ice using RIPA buffer (Sigma Aldrich) supplemented with protease inhibitor cocktail (Sigma), phosphatase inhibitor cocktail (Sigma), sodium orthovanadate (NEB, 1mM), sodium fluoride (NEB, 1 mM), and sodium butyrate (100uM). 150uL of ice cold buffer was added to heads followed by immediate homogenization with a motorized pestle for 10 seconds on ice. Lysates were incubated on ice for 10 minutes followed by 20 seconds of sonication and centrifugation at 16000xg at 4C for 10 minutes. Protein lysates were added 1:1 to 2X protein sample buffer (1mM Tris-HCL pH 6.8, 10% SDS, 1% Bromophenol blue, and 1M DTT) and denatured at 95°C for 10 minutes. Protein was separated by SDS-PAGE on a 4-12% gel (Biorad) at 200V for 30 minutes, followed by electrophoretic transfer to PVDF or nitrocellulose membrane at 70V for 1 hour. Blots were incubated in 5% milk in .1% TBS-T at room temperature for one hour, followed by overnight incubation with primary antibodies overnight at 4°C. Membranes were washed with .1% TBS-T and incubated with HRP-conjugated secondary antibodies (abcam) at room temperature for 1-4 hours. Membranes were washed again with .1% TBS-T and then incubated briefly in ECL substrate (SuperSignal West Femto, ThermoFisher) before imaging. Band detection and quantification was performed using Image Lab software (Bio-Rad). Rabbit anti-histone H3 (abcam, ab1791, 1:20000), Rat anti-histone H3K9ac (active motif, #61663, 1:500), Rabbit anti-LaminB1(CST, #13435S, 1:2000) Rabbit anti-Beta-Tubulin (abcam, ab179513, 1:500), rabbit anti-pS6K(T398) (#9209S 082813, 1:1000), mouse anti-H4K9me3 (Millipore #05-1242, 1:1000), rabbit anti-H3K27ac (D. Lombard, 1:2000), rabbit anti-ubiquitin (Santa Cruz 1:5000), Rabbit anti-histone H4 (abcam, ab10158 1:5000), mouse anti-GAPDH (proteintech #60004 1:1000), rabbit anti-dS6K (T. Neufeld 1:3500), rabbit anti-pAKT (CST 1:1000), rabbit anti-AKT (CST 1:1000), and rabbit anti-GFP (abcam, 1:1000) were used for primary antibody staining.

### RNA extraction and RT-qPCR

Experimental flies were flash frozen in liquid nitrogen after appropriate diet exposures. Heads and bodies were separated using a metal sieve on dry ice, and 10 heads were pooled for each biological replicate. Total RNA was extracted using TRIzol Reagent (Thermo Fisher Scientific, #15596026) following the manufacturer’s instructions. Flies were collected into nuclease-free lysing tubes with matrix D beads (MPbio, #6913-500-129984) and 300 ul of TRIzol, then homogenized for two 15-s pulses at 6.5 M/s. Lysates were incubated at room temperature for 10 min, and then 100 ul of chloroform (Sigma-Aldrich) was added for phase separation. 175 ul of RNA containing supernatant was transferred to nuclease-free tubes and mixed with an equal volume of isopropanol (Sigma-Aldrich) to precipitate the RNA, and centrifuged at 12000xg for 15 min at 4°C. The pellet was washed twice with 70% cold EtOH (Sigma-Aldrich), air dried, and redissolved in 25 ul of nuclease-free water. RNA concentration and quality was determined using a NanoDrop One (Thermo Fisher Scientific). Complementary DNA (cDNA) was prepared from 1 ug of total RNA using High-Capacity cDNA Reverse Transcription Kit (Thermo Fisher Scientific) and the resulting cDNA was diluted 1:50 before use. No-RT reactions were prepared and run as controls to ensure no genomic DNA contamination. Quantitative polymerase chain reactions (qPCRs) contained 1X PowerUp SYBR Green PCR Master Mix (Applied Biosystems), 500 nM appropriate primers, and 5 ul 1:50 cDNA for a total reaction volume of 10 ul. C^T^ values were calculated using an absolute threshold of 1′Rn=0.1 and relative expression was determined using the comparative C^T^ approach ^66^. Primers used in this study are:

*RpL-32-RA*_Forward (cgg atc gat atg cta agc tgt) and
*RpL-32-RA*_Reverse (gcc ctt gtt cga tcc gta)
*H3.3A*_Forward (GAAGAAGCCACATCGCTACC) and
*H3.3A*_Reverse (CACAGATTGGTGTCCTCGAA)
*H3*_Forward (ACCGAGCTTCTAATCCGCAAG) and
*H3*_Reverse (ACCAACCAGGTAGGCTTCGC)
*CG1673*_Forward (TGCGCTTTTACTTCCAAGCAGCA) and
*CG1673*_Reverse(GGGCCTAGGTTCTACTGACGGGT)
*Trh*_Forward(GTGCTCCAGTTTTGACTTCGG) and
*Trh*_Reverse(TTTACGGTACACGGGGTCCT)
*Tph*_Forward(CCTCTGTACTATGTGGCCGA) and
*Tph*_Reverse(TCGAGTCGAGAACCTCAACA)

### Brain Immunohistochemistry

Adult brain immunostaining was performed as previously described^67^. Adult brains were dissected and fixed in PBS, pH 7.4 containing 3.7% formaldehyde for approximately 1 hr. Fixed brains were washed quickly 3 times followed by 3 – 20 min washes in 0.1% PBS-T, or 0.5% PBS-T for histone immunostaining, with gentle shaking at room temperature. Brains were blocked using 5% normal goat serum (NGS) in 0.1% PBS-T for at least 30 min at room temperature with gentle rocking, then incubated in primary antibody diluted in 5% NGS for two nights at 4°C. After primary antibody incubation, brains were washed 3 times in PBS-T and incubated in secondary antibody diluted 1:500 in 5% NGS for one night at 4°C. Brains were washed 3 times in PBS-T and mounted between a glass microscope slide and a #1.5 cover glass separated by a custom bridge in VECTASHIELD Antifade Mounting Medium (Vector Laboratories). Samples were imaged on a Nikon A1 Confocal Microscope using either a 20X air or 40X oil lens objective. All treatments were mounted under the same cover slip and at least two slides per experiment were imaged. Image processing was performed using ImageJ (NIH). ROIs were drawn by hand around appropriate cell bodies and values are background-subtracted. Images are representative maximum intensity projections compiled from 1-2 μm thick sections of the indicated number of z-stacks and are contrast matched. Rabbit anti-serotonin (Sigma, S5545 1:6000), mouse anti-nc82 (DSHB 1:20) and rabbit anti-histone H3 (abcam, ab1791 1:20000) were used for primary antibody staining, and Alexa Fluor 488 and 594 were used for secondary antibody staining (Life Technologies 1:1000).

### Egg laying assay

Following eclosion, male and female flies were allowed to mate on SY10% food for 48 hours. Groups of 5 males and 5 females were sorted to appropriate holidic diets. Food was changed every day for the first 2 days and e/o day thereafter and eggs were counted from the old media. Egg counts were obtained from 8-10 vials per treatment.

### Triacylglyceride and Total Protein Quantification

Experimental flies were flash frozen in liquid nitrogen after exposure to appropriate diets for one week. Two experimental flies per biological replicate were homogenized in 200 μl PBS/0.05% Triton-X with protease inhibitor cocktail (Sigma). The homogenate was added into Infinity Triglyceride Reagent (Thermo Electron Corp.) according to manufacturer’s instructions. In parallel, the homogenate was added to BCA working reagent (Pierce BCA Protein Assay Kit) according to manufacturer’s instructions. TAG concentrations were determined by the absorbance at 520nm and estimated by a known triglyceride standard. Total protein concentrations were determined by absorbance at 562 nm and estimated by a known protein albumin standard.

### Activity Assay

Activity recordings and data processing were performed using the *Drosophila* Activity Monitor System (TriKinetics). After diet exposure for 5 days, adult flies were individually tested in 5 mm x 65 mm polycarbonate tubes with the appropriate diet food at one end of the testing tube. The first day of data was removed from the final analysis in order to allow for acclimation to experimental housing conditions. Total activity counts were calculated for each fly by summing all activity counts recorded during the light and dark cycle, respectively. Experiments were performed at 25°C and 60% humidity under a 12-hour light:12-hour dark cycle.

### Statistics

Pairwise comparisons between treatment survivorship curves were carried out using the statistical package R with DLife, as previously described^65^. P-values were obtained using the log-rank test. For all other comparisons involving only one level, we used Student’s t-test to detect significant differences between two treatments or one-way ANOVA followed by Tukey’s post-hoc test after verifying normality and equality of variances. T-tests were two-tailed during initial characterization experiments, or one-tailed in future experiments where the predicted direction of change was known. For comparisons involving more than one level, we used two-way ANOVA to detect significant interactions between the levels and followed up with Tukey’s post-hoc when significance was detected (p<0.05). In cases where data were non-normally distributed (FLIC data), we performed a box-cox transformation to the 0.25 power before computing P-values. In cases where experimental replicates were pooled, a two-way ANOVA with blocking for experiment was performed to ensure non-significant experimental effects. P-values for experiments with less than 3 biological replicates per treatment are not reported. For all dot and bar plots, error bars represent the SEM. All statistical tests and graphing were performed using R. Specific details of statistical analyses are presented in the figure legends.

## Acknowledgments

We wish to thank past and present members of the Pletcher laboratory for their support and comments about the experimental design of these studies, and express gratitude to David Paris for his help engineering and creating some tools used in this study. We also thank members of the Lombard laboratory for sharing antibodies and protocols. We acknowledge Binyamin Jacobovitz in the Michigan Medicine Microscopy Core for training and advice on confocal imaging. **Funding:** This research was supported by The National Science Foundation Graduate Research Fellowship Program (No. DGE 1256260) and the Howard Hughes Medical Institute through the James H. Gilliam Fellowships for Advanced Study program to K.J.W (#GT11426) and the US National Institute of Health, National Institute on Aging (R01 AG051649, R01 AG030593, and R01 AG063371) and the Glenn Medical Foundation to S.D.P.

## Author contributions

Conceptualization: KJW, SDP

Methodology: KJW, SDP

Investigation: KJW, RAH, EH, SDP

Visualization: KJW, EH

Funding acquisition: KJW, SDP

Project administration: KJW, SDP

Supervision: KJW, SDP

Writing – original draft: KJW, SDP

Writing – review & editing: KJW, SDP

## Supplementary Materials for

**Figure S1.**
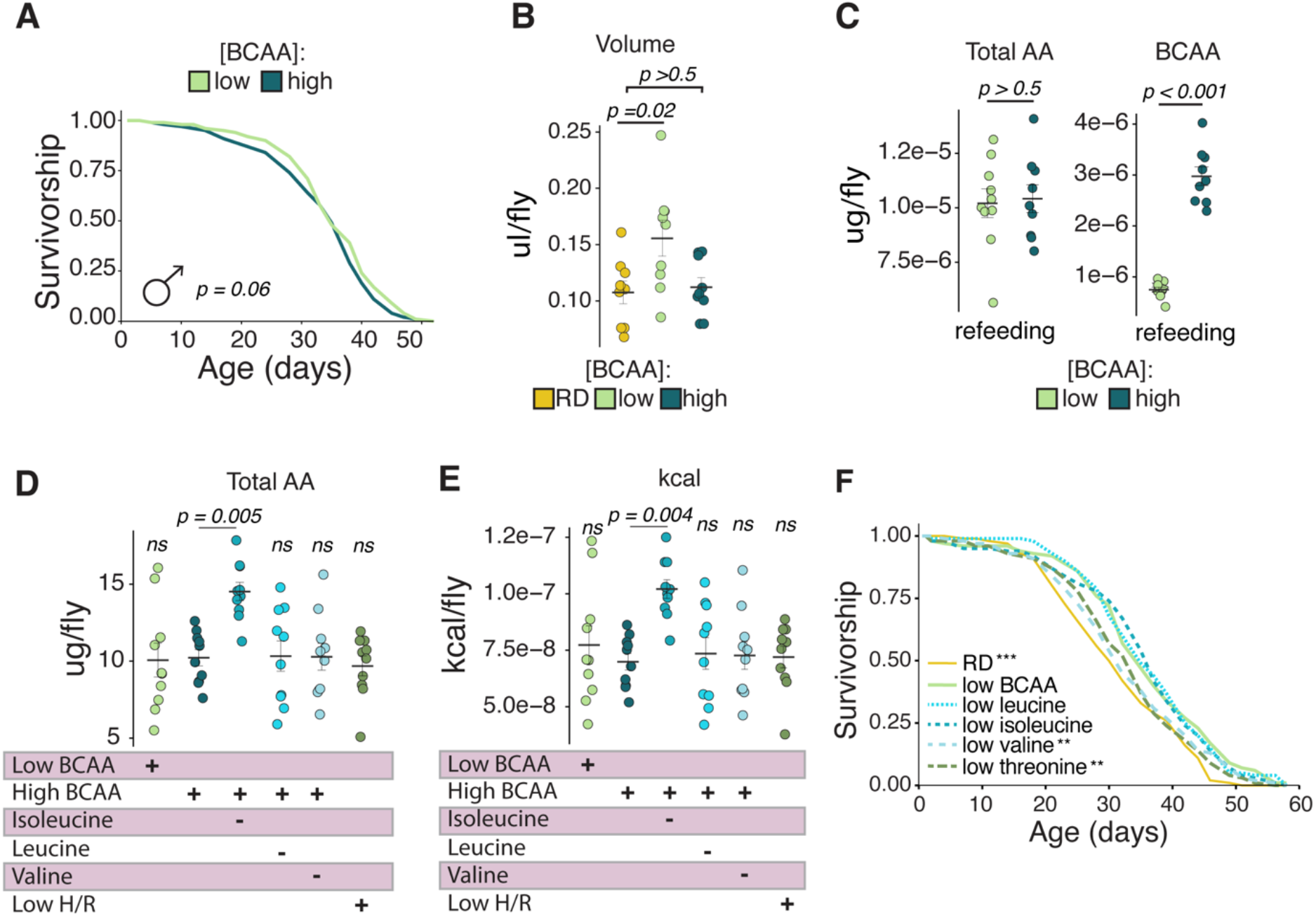
Isoleucine is necessary and sufficient to increase feeding and modulate aging. **(A)** Lifespan of male *Canton-S* flies on low- or high-BCAA diets, related to Figure 1B (log-rank test, N= 217-231**). (B)** Total volume consumed by female *Canton-S* flies on RD, low-, or high-BCAA diets for 1 week (one-way ANOVA with Tukey’s post-hoc). **(C)** Total amino acid and BCAA consumed during 3-hour refeeding low- or high-BCAA food, related to Figure 1I (one-way ANOVA). **(D-E)** Total amino acid **(D)** and kcal **(E)** consumed by *Canton-S* flies on indicated diets, related to Figure 1J. **(F)** Lifespans of *Canton-S* flies on indicated diets (log-rank test, p-values derived by comparison to low-BCAA, N=170-177, **p<0.01, ***p<0.001).

**Figure S2.**
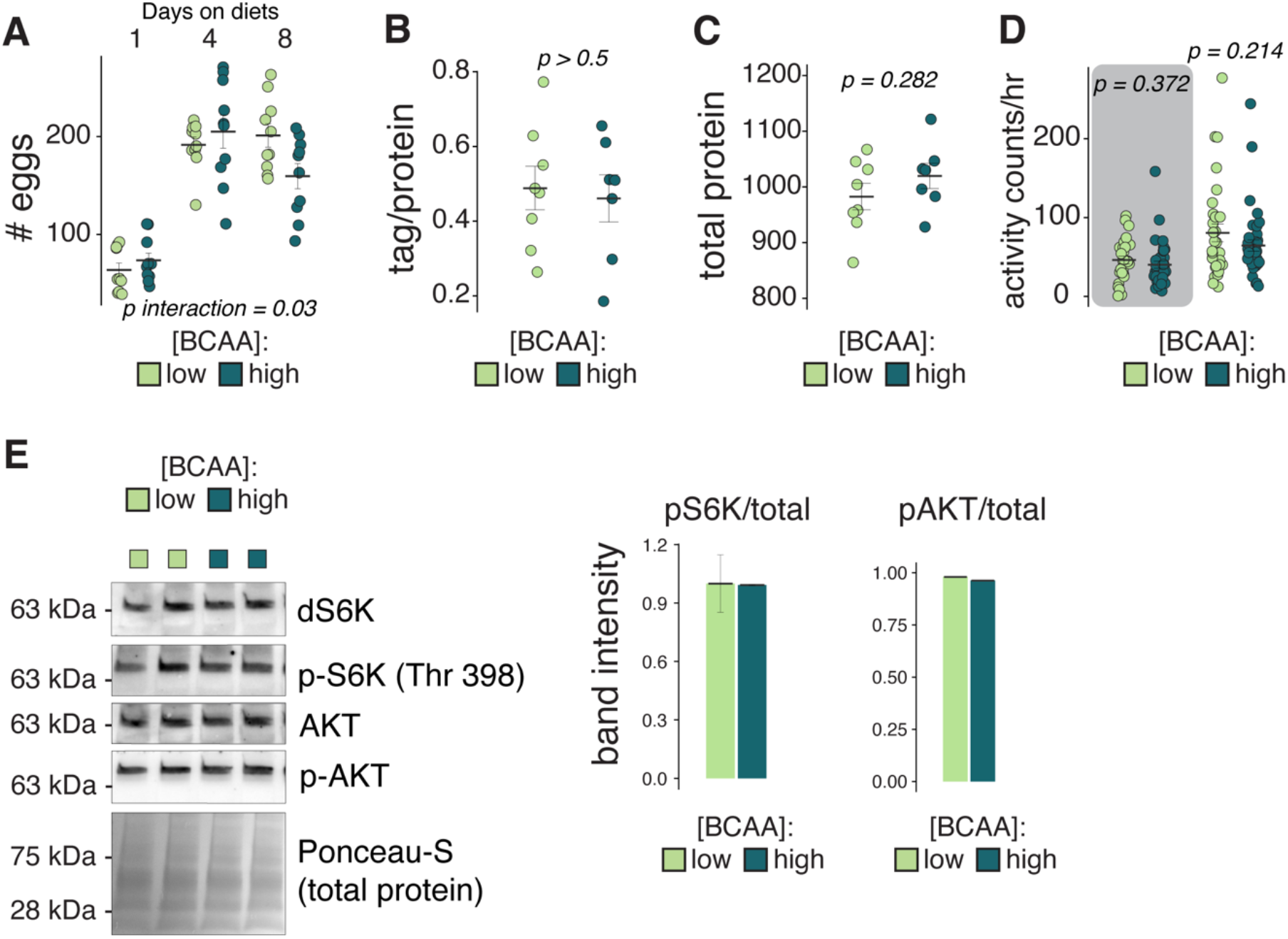
Physiological measures typically associated with aging are unchanged by dietary BCAAs. **(A)** Eggs laid by *Canton-S* flies on indicated diets after 1, 4, or 8 days on diet (two-way ANOVA). **(B)** Total triacylglycerides in *Canton-S* flies after 1 week on BCAA diets (one-way ANOVA, 2 flies per biological replicate). **(C)** Total protein in *Canton-S* flies after 1 week on BCAA diets (one-way ANOVA, 2 flies per biological replicate). **(D)** Total activity counts of *Canton-S* flies after 5 days on BCAA diets during dark (left) and light (right) period (one-way ANOVAs). **(E)** Western blot and quantification of S6K and AKT activation in heads of *Canton-S* flies after 1 week on BCAA diets (N=2).

**Figure S3.**
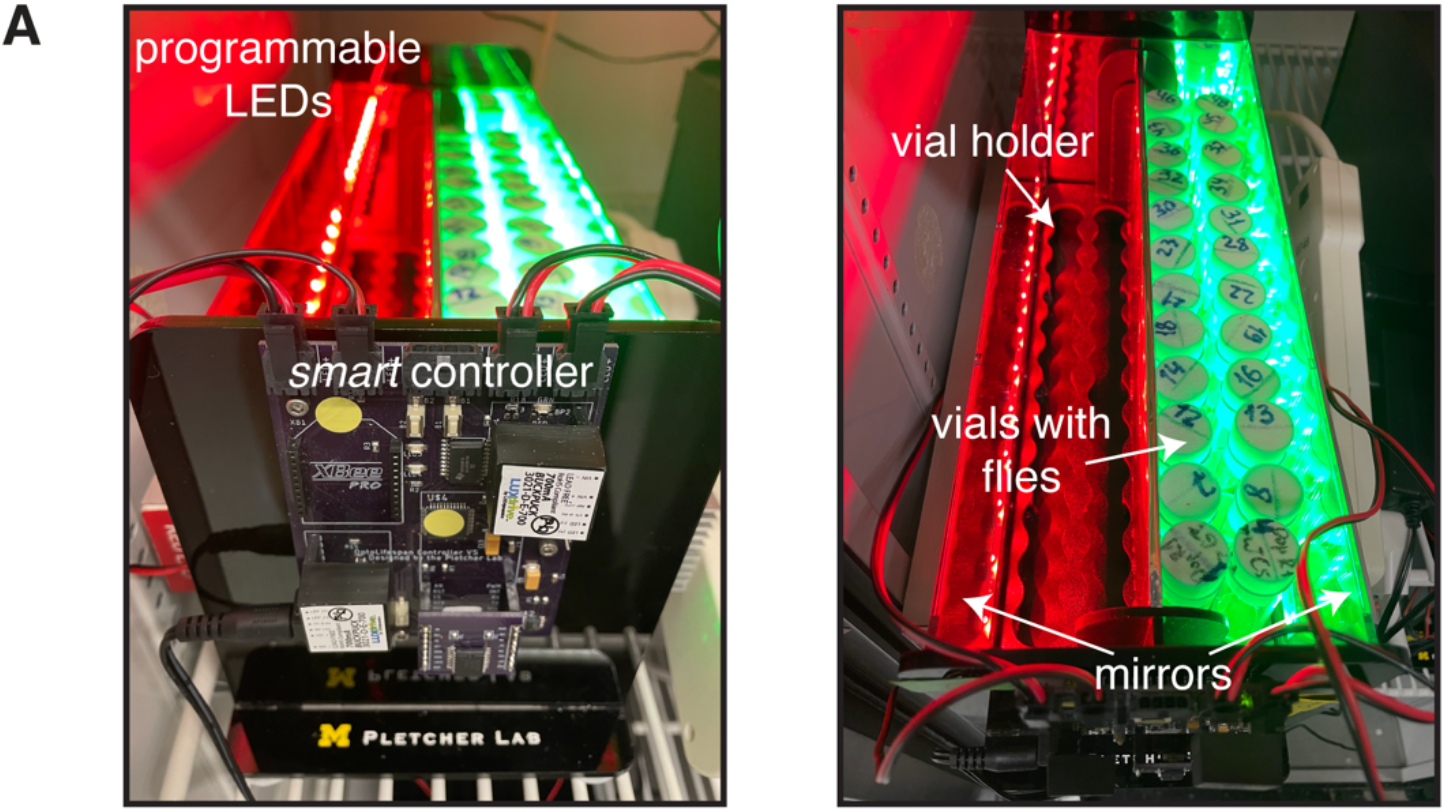
Custom rig for optogenetic experiments. **(A)** Representative rig used for optogenetic lifespan and Con-Ex experiments. Details can be found in methods.

**Figure S4.**
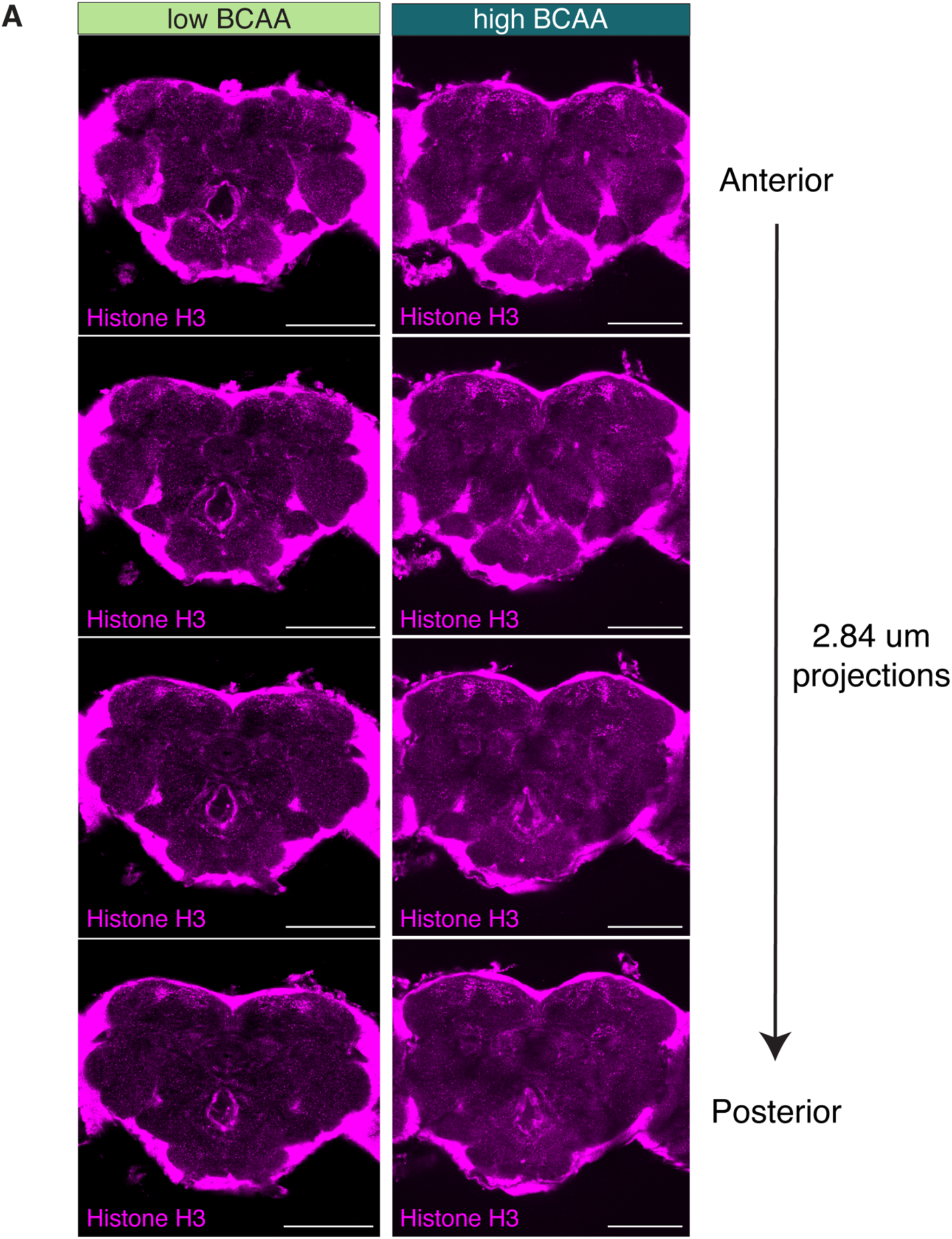
Histone H3 abundance is modulated by dietary BCAAs in discrete anatomical locations. **(A)** Immunostaining for histone H3 in *Canton-S* flies after 5-7 days on BCAA diets. Representative images are montages of maximum intensity projections through the central brain, each consisting of two 1.42μm stacks (scale bar = 100μm).

**Figure S5.**
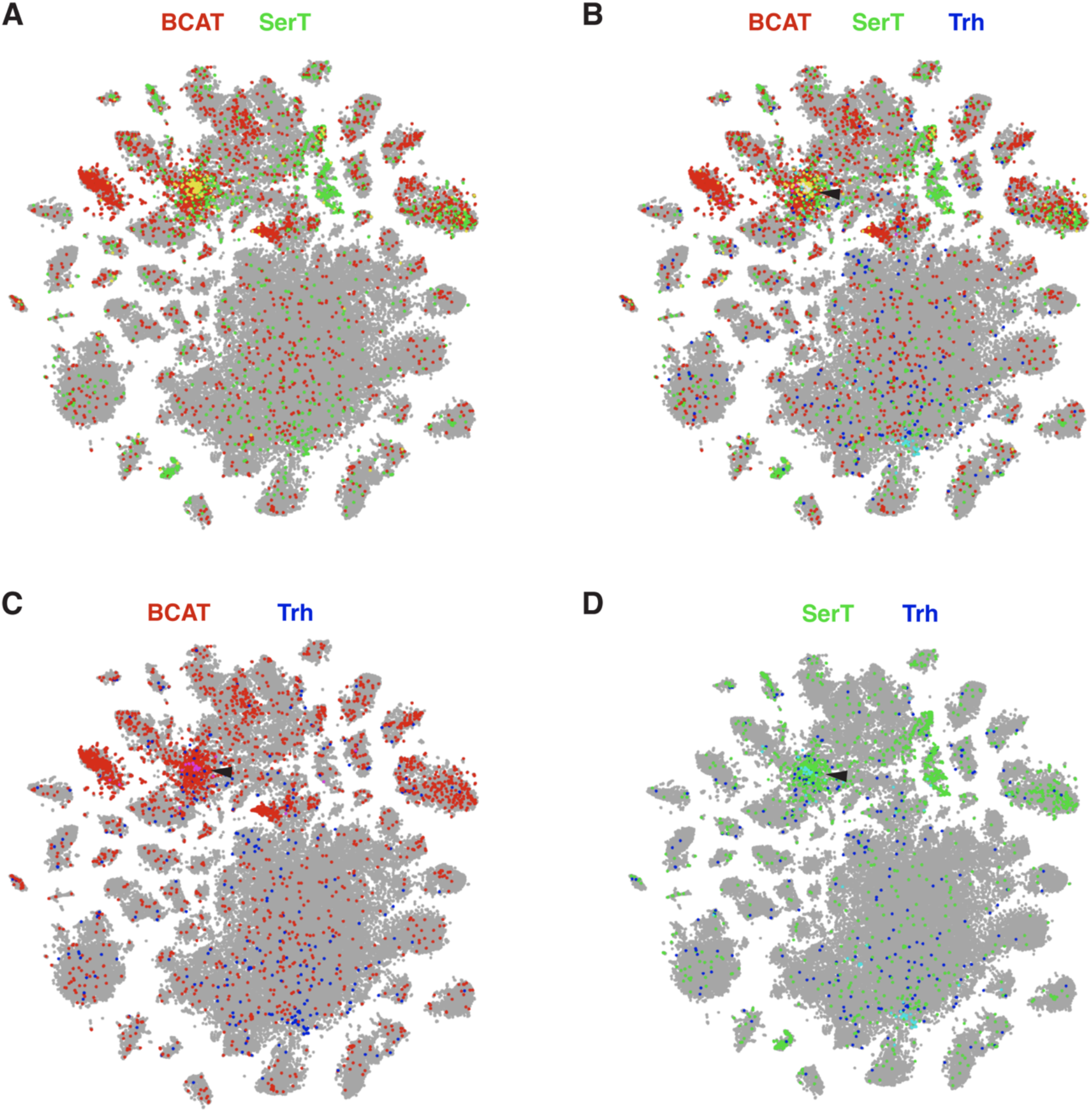
tSNE plots of single-cell gene expression in fly heads. **(A-D)** tSNE plots from the Fly Cell Atlas (www.flycellatlas.org) generated using the publicly available 10x droplet based single-cell sequencing dataset from fly heads and visualized using SCope^47^. **(A)** Cells that express *BCAT* (red) and *SerT* (green) with a cluster of co-expressing cells shown in yellow (approximately 80 cells were detected to express both *BCAT* and *SerT*). **(B)** Cells that express *BCAT* (red), *SerT* (green), *Trh* (blue), and co-expressing cells shown in white and highlighted with arrowhead. **(C)** Cells that express *BCAT* (red) or *Trh* (blue) and co-expressing cells shown in pink. **(D)** Cells that express *SerT* (green) or *Trh* (dark blue) and co-expressing cells shown in light blue.

**Figure S6.**
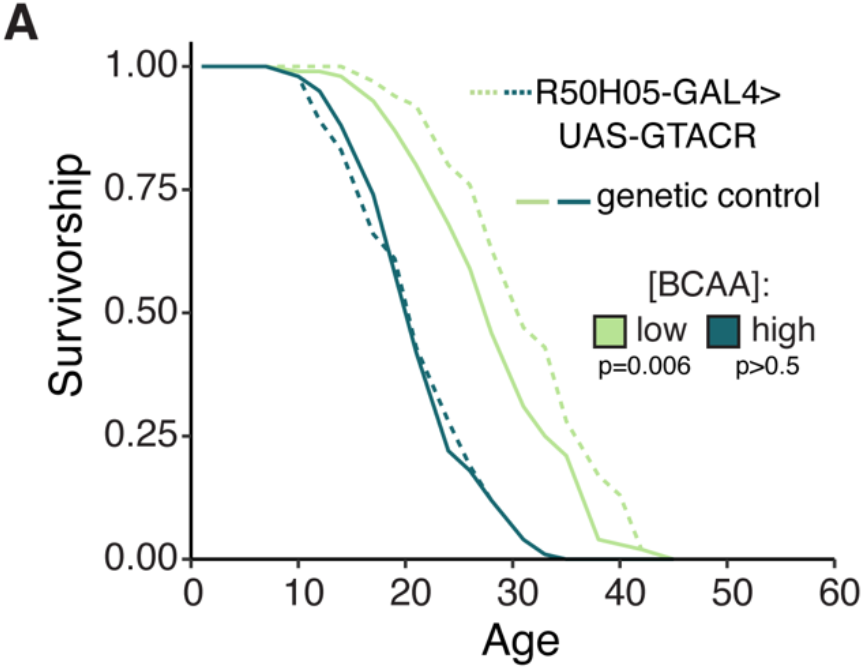
Lifespan of flies with optogenetic inhibition of R50 hunger neurons. Flies carrying *R50H05-GAL4>UAS-GTACR* or *UAS-GTACR/w-;CS* as control were aged on low- or high-BCAA diets and exposed to 530nm light constantly for the duration of the experiment (log-rank test, N=93-100).

**Figure S7.**
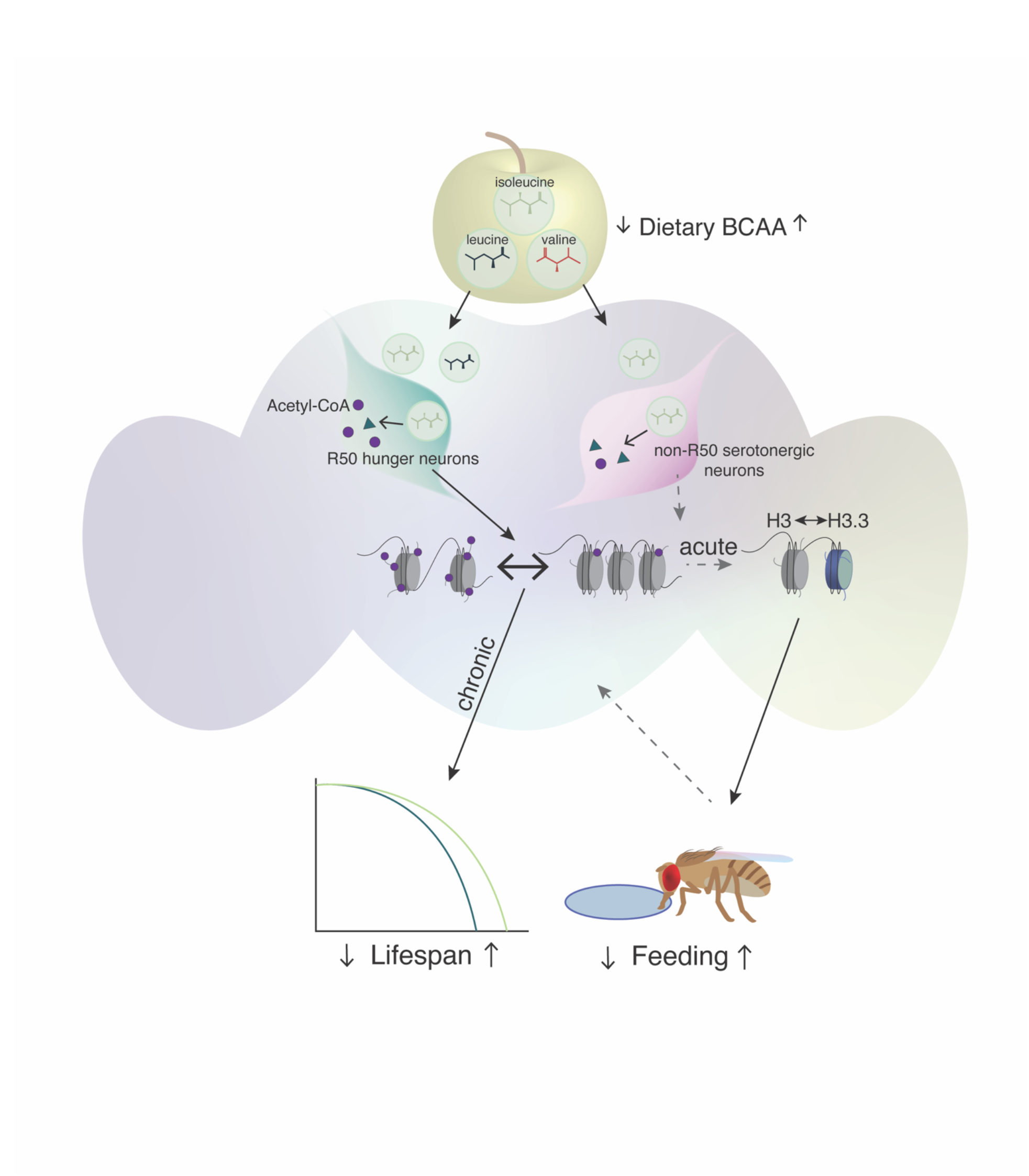
Proposed model. The motivational state of hunger modulates feeding and aging via distinct pathways.

**Table S1.** Composition of diets used for experiments.

| Diet/<br>Component | Yaa<br>(Piper, et<br>al.) | RD | Low<br>BCAA | High BCAA | High BCAA<br>–Isoleucine | High BCAA<br>– Leucine | High BCAA<br>– Valine | Low BCAA<br>+ Isoleucine | Low Histidine/<br>Arginine |
| --- | --- | --- | --- | --- | --- | --- | --- | --- | --- |
| Total volume (l) | 1 | 1 | 1 | 1 | 1 | 1 | 1 | 1 | 1 |
| Agar (g) | 7 | 11.25 | 11.25 | 11.25 | 11.25 | 11.25 | 11.25 | 11.25 | 11.25 |
| Isoleucine (g) | 1.16 | 3.48 | 1.16 | 5.8 | 1.16 | 5.8 | 5.8 | 5.8 | 3.48 |
| Leucine (g) | 1.64 | 4.92 | 1.64 | 8.2 | 8.2 | 1.64 | 8.2 | 1.64 | 4.92 |
| Valine (g) | 1.33 | 3.99 | 1.33 | 6.65 | 6.65 | 6.65 | 1.33 | 1.33 | 3.99 |
| Tyrosine (g) | 0.84 | 2.52 | 2.52 | 2.52 | 2.52 | 2.52 | 2.52 | 2.52 | 2.52 |
| Sucrose (g) | 17.12 | 51.36 | 51.36 | 51.36 | 51.36 | 51.36 | 51.36 | 51.36 | 51.36 |
| Cholesterol<br>solution (ml) | 15 | 15 | 15 | 15 | 15 | 15 | 15 | 15 | 15 |
| CaCl<sub>2</sub> (1000X, ml) | 1 | 1 | 1 | 1 | 1 | 1 | 1 | 1 | 1 |
| MgSO<sub>4</sub> (1000X, ml) | 1 | 1 | 1 | 1 | 1 | 1 | 1 | 1 | 1 |
| CuSO<sub>4</sub> (1000X, ml) | 1 | 1 | 1 | 1 | 1 | 1 | 1 | 1 | 1 |
| FeSO<sub>4</sub> (1000X, ml) | 1 | 1 | 1 | 1 | 1 | 1 | 1 | 1 | 1 |
| ZnSO<sub>4</sub> (1000X, ml) | 1 | 1 | 1 | 1 | 1 | 1 | 1 | 1 | 1 |
| Acetate buffer<br>(10X, ml) | 100 | 100 | 100 | 100 | 100 | 100 | 100 | 100 | 100 |
| Nucleic acids/lipids<br>(125X, ml) | 8 | 8 | 8 | 8 | 8 | 8 | 8 | 8 | 8 |
| EAA solution (ml) | 60.51 | 181.5 | 181.5 | 181.5 | 181.5 | 181.5 | 181.5 | 181.5 | 181.5 |
| NEAA solution (ml) | 60.51 | 181.5 | 181.5 | 181.5 | 181.5 | 181.5 | 181.5 | 181.5 | 181.5 |
| Glutamate solution<br>(ml) | 18.21 | 54.5 | 54.5 | 54.5 | 54.5 | 54.5 | 54.5 | 54.5 | 54.5 |
| Cysteine solution<br>(ml) | 5.28 | 15.5 | 15.5 | 15.5 | 15.5 | 15.5 | 15.5 | 15.5 | 15.5 |
| Vitamin solution<br>(47.6X, ml) | 21 | 21 | 21 | 21 | 21 | 21 | 21 | 21 | 21 |
| Folic acid (1000X,<br>ml) | 1 | 1 | 1 | 1 | 1 | 1 | 1 | 1 | 1 |
| Propionic acid (ml) | 6 | 6 | 6 | 6 | 6 | 6 | 6 | 6 | 6 |
| Tegosept (ml) | 15 | 15 | 15 | 15 | 15 | 15 | 15 | 15 | 15 |

**Table S2.**
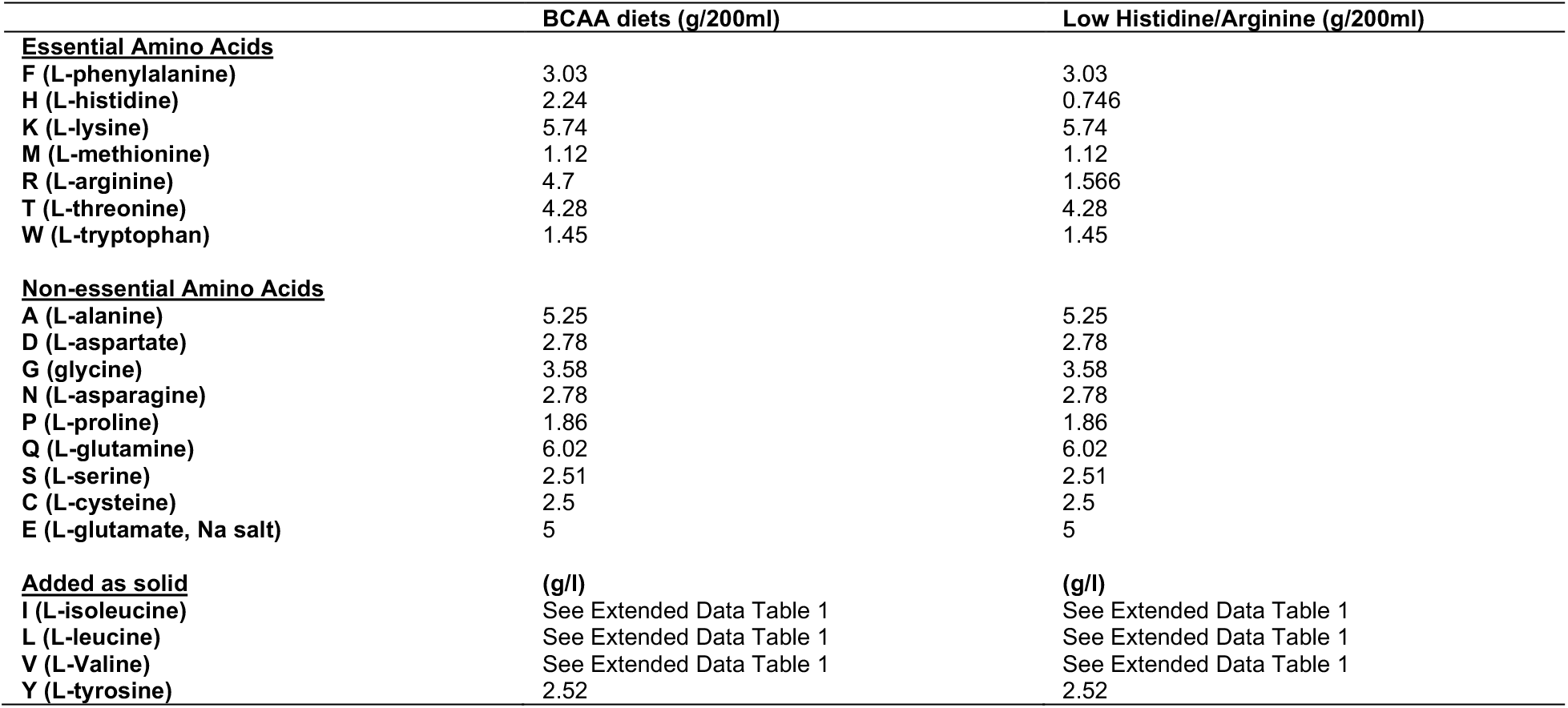
Concentration of amino acid stock solutions used in diets.

